# Sweet chirality: the taste of l- and d-glucose stereoisomers

**DOI:** 10.1101/2020.08.16.252718

**Authors:** Nitzan Dubovski, Yaron Ben Shoshan-Galeczki, Einav Malach, Masha Y Niv

**Affiliations:** The Institute of Biochemistry, Food Science and Nutrition, Robert H. Smith Faculty of Agriculture, Food and Environment, The Hebrew University of Jerusalem, P.O. Box 12, Rehovot 76100, and Fritz Haber Center for Molecular Dynamics, The Hebrew University, Jerusalem, Israel

## Abstract

Chirality plays a key role in biomolecular recognition. Typically, a change in chirality will dramatically affect ligand–receptor binding. However, both d-glucose and its enantiomer l-glucose elicit sweet taste in humans. We show that l- and d-glucose are perceived as similarly sweet by humans, and that in cell-based functional assays, both enantiomers activate the human sweet taste receptor TAS1R2/TAS1R3. We hypothesize that both l- and d-glucose occupy the orthosteric binding site in the VFT domain of TAS1R2. Using induced-fit docking to a homology model of this domain, we identify two subpockets in this binding site. The model suggests that glucose molecules can bind in either of these subpockets, which overlap with the predicted positions of monosaccharide units of sucrose. One subpocket is close to the hinge between the two lobes, and overlaps with aspartame and neotame site, the second subpocket overlaps with that described for sweetness enhancers. These findings suggest a framework for rational design of sweeteners combinations.

## Introduction

Chirality is a fundamental property of molecular asymmetry. One of the aspects of chirality is stereosensitivity, which is based on chiral discrimination (Ariens, 1984). Stereosensitivity is a hallmark of ligand–receptor recognition, due to the specific and complementary bonds between the ligand and the receptor binding site. A chiral center is a tetrahedral atom that has four different substituents. A molecule with a single chiral center has two enantiomeric forms, which are mirror images of each other. Key molecules in nature, such as carbohydrates, amino acids, and nucleic acids, harbor chiral centers. Natural amino acids have l-chirality, whereas sugars have d-chirality, and living organisms typically use only one form of chiral molecules (Blackmond, 2010). This, in turn, dictates the secondary structure of proteins and gives rise to handedness in DNA helices (Wing et al., 1980).

Enantiomers of many molecules can differ in odor quality and intensity, as perceived by humans (Brookes, Horsfield, & Stoneham, 2009). One of the best-known examples of this phenomenon are the carvone enantiomers: (4/*R*)-(–)-carvone smells minty, whereas (4S)-(+)-carvone smells like caraway (Russell & Hills, 1971). The observed variation between enantiomers in terms of odor quality and intensity is attributed to chiral selectivity, causing the enantiomers to access or activate the receptor in different ways (Brookes, Horsfield, & Stoneham, 2009).

The sense of taste is essential for food choice and consumption (Chaudhari & Roper, 2010). Sweet taste is critical for detecting high-calorie foods and essential nutrients, which maintain glucose levels in the body. Preferences for different taste modalities have a strong innate component: sweet, umami (amino acid), and salty substances are innately preferred, whereas bitter and many sour substances are innately rejected (Steiner, Glaser, Hawilo, & Berridge, 2001).

Whereas most naturally occurring l-amino acids taste bitter (Bachmanov et al., 2016; Kawai, Sekine-Hayakawa, Okiyama, & Ninomiya, 2012; Susan S. Schiffman, Clark, & Gagnon, 1982), and most (synthetic) d-amino acids taste sweet (Bassoli, Borgonovo, Caremoli, & Mancuso, 2014; G. E. DuBois, 2016), both stereoisomers of glucose have been reported to have similar taste quality and intensity (B. Dubois, Padovani, Scheltens, Rossi, & Dell’Agnello, 2016; Shallenberger & Acree, 1967). Interestingly, the natural substrate for glycolysis is d-glucose and not l-glucose (Cox & Nelson, 2000), as demonstrated by the latter’s low energy values (Levin, Zehner, Saunders, & Beadle, 1995). l-glucose is also non-transportable; its paracellular movement occurs via diffusion through tight junctions, down the concentration gradient (Kalsi et al., 2009).

Recognition of the sweet taste modality is mediated by sets of chemosensory guanine nucleotide-binding protein (G protein)-coupled receptors (GPCRs) expressed in separate taste cells on the tongue (Chaudhari & Roper, 2010). Sweet compounds are primarily detected by a heterodimer consisting of two subtypes of type 1 taste receptor (TAS1Rs): TAS1R2 and TAS1R3, both belonging to the C family of GPCRs (Damak et al., 2003; Sclafani, Zukerman, & Ackroff, 2020; Zhao et al., 2003). Family C GPCRs are homo- and heterodimers, and possess a large extracellular domain that contains the Venus flytrap module where the orthosteric binding site resides, and a cysteine-rich domain. The most studied members of Family C GPCRs are the metabotropic glutamate receptor (mGluR) subtypes, and the gamma-aminobutyric acid (GABA) receptor. Several structures of the mGluRs (Christopher et al., 2019; Schkeryantz et al., 2018; D. Zhang, Zhao, & Wu, 2015) and GABA_B_ receptor (Mao et al., 2020; Papasergi-Scott et al., 2020) have been solved. To date, no experimental structure of human taste receptors exists. Defining the structural mechanisms used by these GPCRs to detect the diversity of compounds, while simultaneously achieving high selectivity for specific types of molecules, remains a challenge. However, structural information on the sweet taste receptor is available for the medaka fish: recently, Nuemket et al. determined the X-ray structure of the medaka fish TAS1R2-N-terminal domain/TAS1R3-N-terminal domain heterodimer (Nuemket et al., 2017). The experimental structure was used as a template for homology modeling of the sweet taste receptor, which in turn was used for docking and binding-site analyses (Abu et al., 2020; Ben Shoshan-Galeczki & Niv, 2020; Di Pizio, Ben Shoshan-Galeczki, Hayes, & Niv, 2019).

Activation of the sweet taste receptor produces modulation of the signal-transduction pathway, through coupling with G proteins. G proteins are complexes that consist of alpha (α), beta (β) and gamma (γ) subunits (Simon, Strathmann, & Gautam, 1991). When a sweet agonist binds to the Tas1R2/Tas1R3 heterodimer, located on taste receptor cells, a second-messenger cascade is activated, through coupling with Gαq. First, phospholipase C β2 is activated, which then stimulates the second messengers inositol triphosphate (IP3) and diacylglycerol. IP3 causes the release of Ca^2+^ from intracellular stores. The calcium opens non-selective TRPM5 channels, which allow cations to enter the cell; this causes depolarization, which generates the perception of sweet taste in the brain (Breer, Boekhoff, & Tareilus, 1990; McCaughey, 2008). The extent of ligand-mediated Gαq-coupled receptor activation can be quantitatively determined by measuring either the release of intracellular Ca^2+^ or the concentration of intracellular IP3. The current study uses a direct measurement of IP3, as detailed in the Methods and following our previous work on bitter (Di Pizio et al., 2020) and sweet taste receptors (Abu et al., 2020).

We first confirm that l-glucose tastes sweet with sensory tests. Then, using the cell-based system, we show that its sweetness is mediated by the TAS1R2/TAS1R3 sweet taste receptor and finally suggest a possible structural explanation for lack of selectivity in this receptor.

## Results

### l-Glucose presents similar sweetness to d-glucose

l-Glucose has been previously reported to have similar taste properties to d-glucose (G. E. DuBois, 2016; Shallenberger, Acree, & Lee, 1969). We confirmed the similarity in sweetness of l-glucose and d-glucose by two-alternative forced choice (2AFC) test: whereas this test showed the expected difference between d-glucose and sucrose (Figure 1A), no significant difference was found between d-glucose and l-glucose with respect to perceived sweetness (Figure 1B).

**Figure 1.**
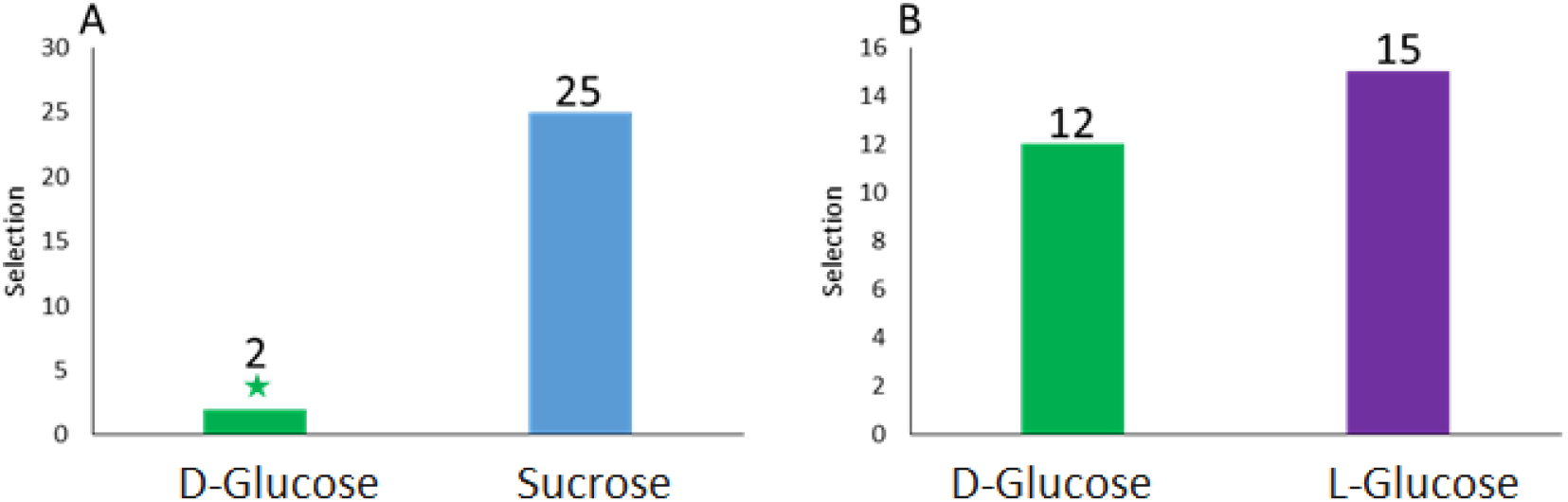
A. Two-alternative forced choice (2AFC) test between 0.48 concentration solutions of sucrose (blue) and d-glucose (green); sucrose is sweeter. B. 2AFC test between d-glucose and l-glucose (purple); there is no significant difference in sweetness

### Individual perception of l- and d-glucose

Next, the sweetness of three concentrations of these sugars was measured using the generalized labeled magnitude scale (gLMS). Two participants perceived d-glucose as sweeter than l-glucose, two other participants perceived l-glucose as sweeter than d-glucose, and two participants perceived both of the monomers as similarly sweet. Although the sweetness of l- and d-glucose might have been perceived slightly differently among individuals, the perception was similar on average. Tukey–Kramer statistical analysis between all six groups of results from all six participants (Figure 2G) showed no significant difference between d-glucose and l-glucose at each concentration, while sweetness increased with increasing concentrations.

**Figure 2.**
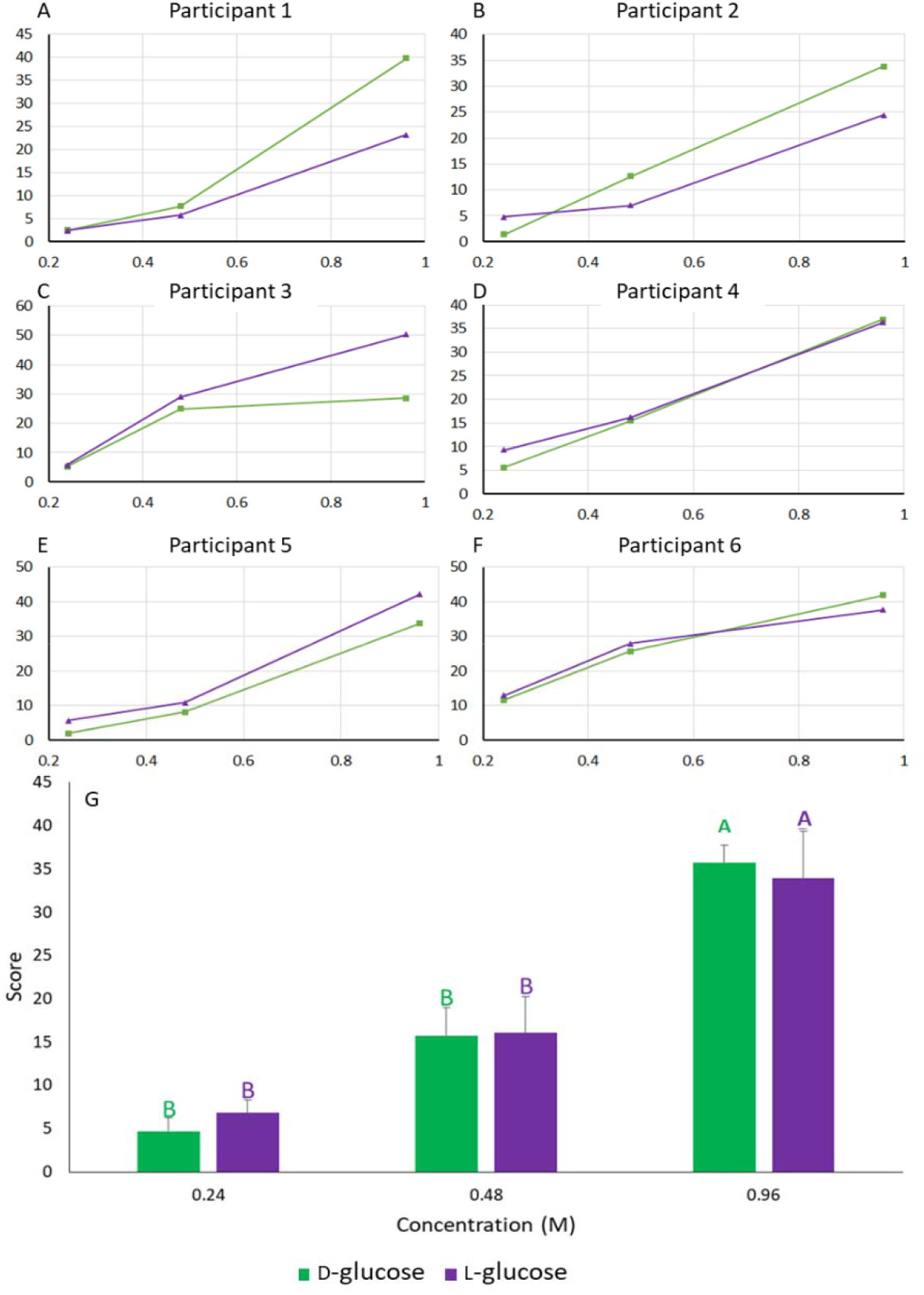
A–F. Labeled magnitude scale (gLMS)—ranging from 0 (barely detectable) to 100 (strongest imaginable)—results for glucose stereoisomers from six trained participants shows similar sweetness of the compounds in a range of concentrations. G. Average scores for the d- and l-glucose solutions given by the six participants across different concentrations.

Upon confirming the sweetness of l-glucose, we tested whether the sweetness of both stereoisomers is mediated by the same receptors, using an in-vitro cell-based assay for human sweet taste receptor TAS1R2/TAS1R3.

#### Characterization of the cell-based system

Carbachol is known to activate the muscarinic (acetylcholine) receptor that is endogenously expressed in HEK293T cells. As a proof of concept for the cell-based assay’s efficiency, we evaluated IP1 levels after treatment with carbachol. Indeed, exposure to 0.00025 M carbachol significantly elevated IP1 levels compared to basal activity (exposure to 0.05 M LiCl) (data not shown). To confirm the expression of the sweet taste receptor in the transiently transfected cell line, the effects of known sugars and sweeteners on IP1 levels were examined. As expected, sucrose, sucralose, and acesulfame-K evoked activation of the sweet taste receptor in a dose-responsive manner (Figure 3).

**Figure 3.**
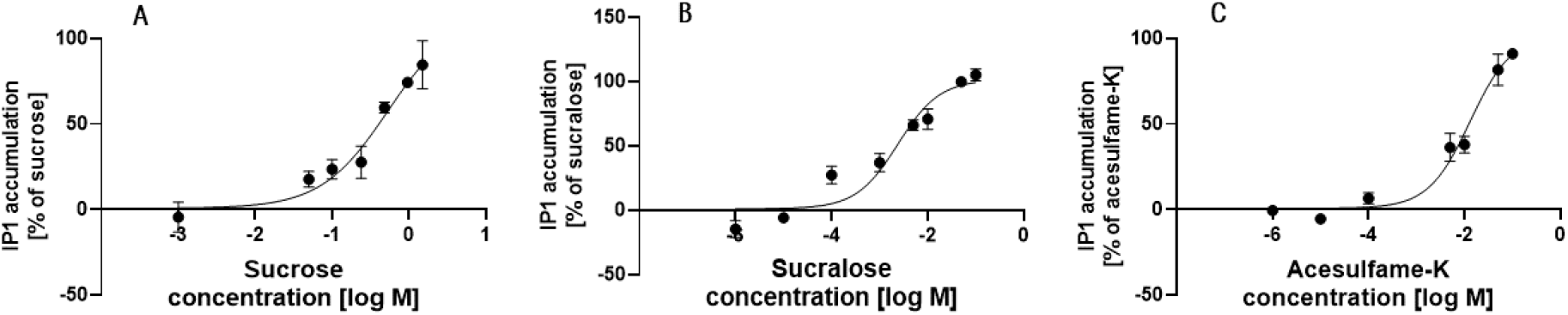
Dose-response curves of HEK293T cells transiently transfected with TAS1R2, TAS1R3 and Gα16gus44, determined by using HTRF IP1-based assay, after application of (A) sucrose, (B) sucralose, and (C) acesulfame-K. Each of the graphs presents the merging of 2–3 independent repetitions.

#### l-Glucose activates the sweet taste receptor

Similar to d-glucose, l-glucose elevated IP1 levels in transfected cells in a dose-responsive manner (Figure 4A-B). In contrast, in nontransfected cells, no difference in IP1 levels was observed between any of the concentrations (see Figure S1). Taken together, these findings support the hypothesis that l-glucose specifically activates the sweet taste receptor.

**Figure 4.**
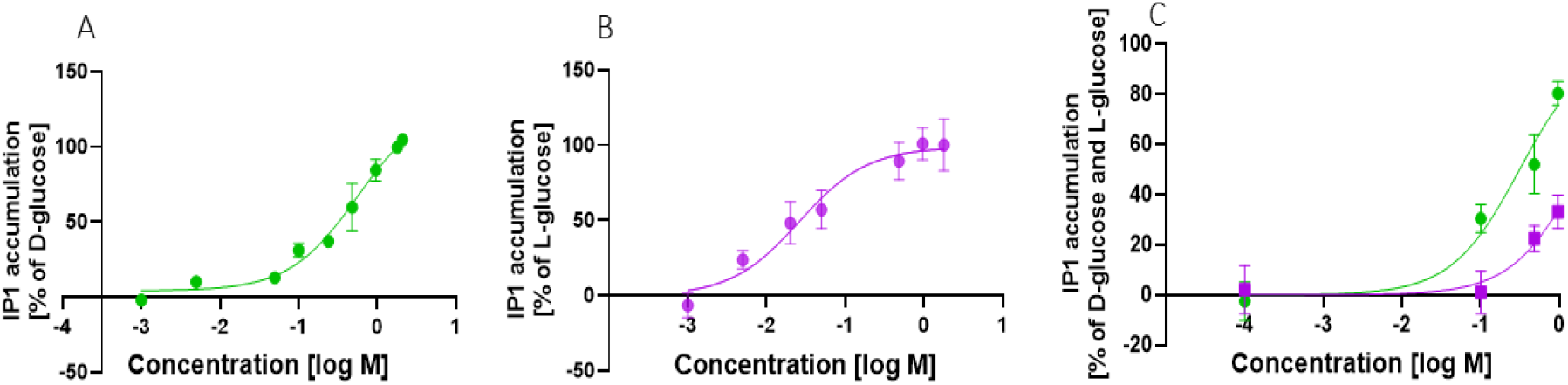
Dose-response curves of HEK293T cells transiently transfected with TAS1R2, TAS1R3 and Gα16gus44, determined by HTRF IP1-based assay, after application with (A) d-glucose and (B) l-glucose, at different concentrations. Each graph (A,B) presents a merging of 3–4 independent repetitions. Graph C illustrates activation by l-glucose (purple line) compared to d-glucose (green line), which was (the latter) used as a reference for the measurement of l-glucose’s level of activation. Graph C represents three independent repetitions which showed the same trend of higher activation by d-glucose than by l-glucose.

Sensory experiments showed that on average, the sweetness of d- and l-glucose is similar (Figure 2G). In contrast, when d-glucose was used as a reference to estimate l-glucose’s level of activation in transfected HEK293T cells, the experiments consistently showed higher activation by d-glucose than l-glucose, as measured by comparison of IP1 levels for each concentration of both stereoisomers (Figure 4C). The inter-individual differences, and a direct comparison between sensory and cell-based results will be explored in future studies.

To understand how the same receptor is activated by different stereoisomers, modeled the interactions between l- and d-glucose and the sweet taste receptor.

### Molecular modeling of sucrose, and l- and d-glucose binding to the receptor

There are several optional conformations for glucose: 5-atom ring (furanose), 6-atom ring (pyranose) and the open form, which does not close to a ring. The distribution of the different isomers in aqueous solution is 1% furanose, 0.002% open form and the rest is present in the the pyranose form (Wittmann, 2006). Thus, the furanose and open forms of glucose account for only up to 1.002% of the isomeric forms, and we focused here solely on the pyranose form of the glucose isomer. d-glucose and l-glucose both have additional conformations as a result of mutarotation, with opening and closing of the rings into different forms, depending on the position of the hydroxyl group on carbon 1: above the plane of the ring (beta) or below the plane of the ring (alpha). Thus, there are four pyranose conformations: α-d-glucose, β-d-glucose, β-l-glucose and α-l-glucose.

### Docking

Modeling of the sweet receptor VFT domain was carried out previously (Abu et al., 2020; Ben Shoshan-Galeczki & Niv, 2020). We use it here to compare the different enantiomers of glucose in the hT1R2 binding site. We performed initial flexible docking of each of the four isomers, followed by induced-fit docking (IFD) as in previous simulations(Di Pizio, Shy, Behrens, Meyerhof, & Niv, 2018).

Docking the three sugars studied here, sucrose, and l- and d-glucose, we found that the two possible pockets of glucose coincide with the pockets occupied by monosaccharide units of sucrose. The results indicated that each monosaccharide can be placed in one of the two subpockets. The first is close to the hinge of the protein between the two binding lobes. The second is adjacent to the first one, but located further away from the hinge of the protein. Thus, modeling suggests that both of the glucose enantiomers bind to the orthosteric binding site in the VFT domain of T1R2, and it is plausible that two glucose molecules bind at the same time (Figure 6).

**Figure 6.**
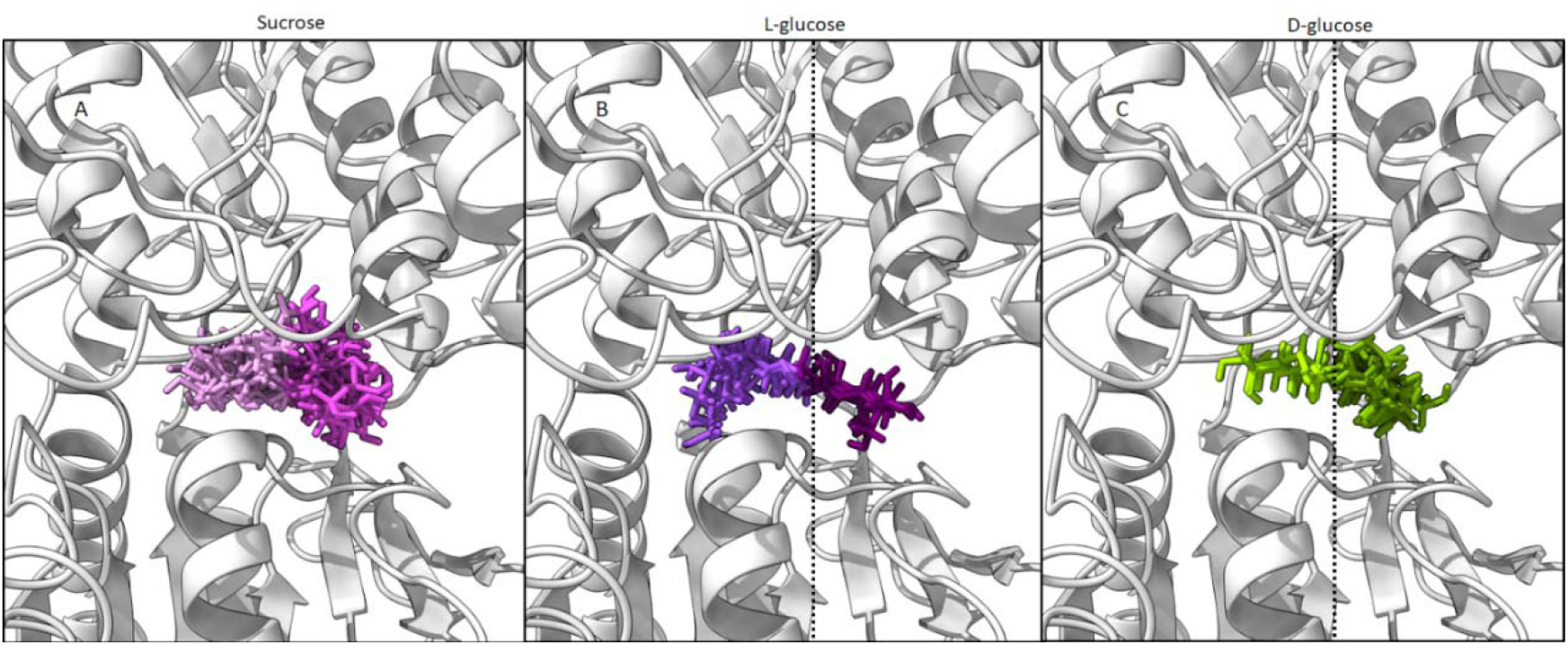
Predicted poses of sucrose (pink), l-glucose (purple) and d-glucose (green). l- and d-glucose cluster in two binding pockets, adjacent to each pocket, while sucrose poses are clustered similarly, such that the two monosaccharides occupy the two binding sites.

Interestingly, d-glucose had more docked poses in the binding pocket that was further away from the hinge, and l-glucose poses were more common in the binding pocket close to the hinge. 2d-interaction analysis was used to identify interacting residues that were within 4 Å from ligands in the binding sites. Some residues were unique to subpocket A (D142, N143 and P277), others to subpocket B (I306, D307 and T326), and some participated in both binding sites. Interestingly, these six residues (seen in Figure 7) have been shown (Maillet et al., 2015; Masuda et al., 2012; F. Zhang et al., 2010) to be involved in the activation of sweet taste receptor by diverse sweet compounds, such as aspartame and neotame. The residues in either one of the subpockets: Y103, D142, N143, S165, Y215, P277, E302, S303, D307, T326, I327 and I382 are conserved in primates and rodents (see Figure 7 and Figure S2). Two exceptions are I67 and D278 that are replaced in mice and rats by residues with similar properties. In mice and rats the I67 is replaced by Leucine, and D278 is replaced by Glutamate. The common residues in both subpockets were mentioned previously as Tas1R2 VFT domain binding residues (Cheron, Golebiowski, Antonczak, & Fiorucci, 2017), while residue D307, which is unique to binding pocket B, was also reported as a unique interacting residue for the sweetness enhancer - SE-3 (F. Zhang et al., 2010).

**Figure 7.**
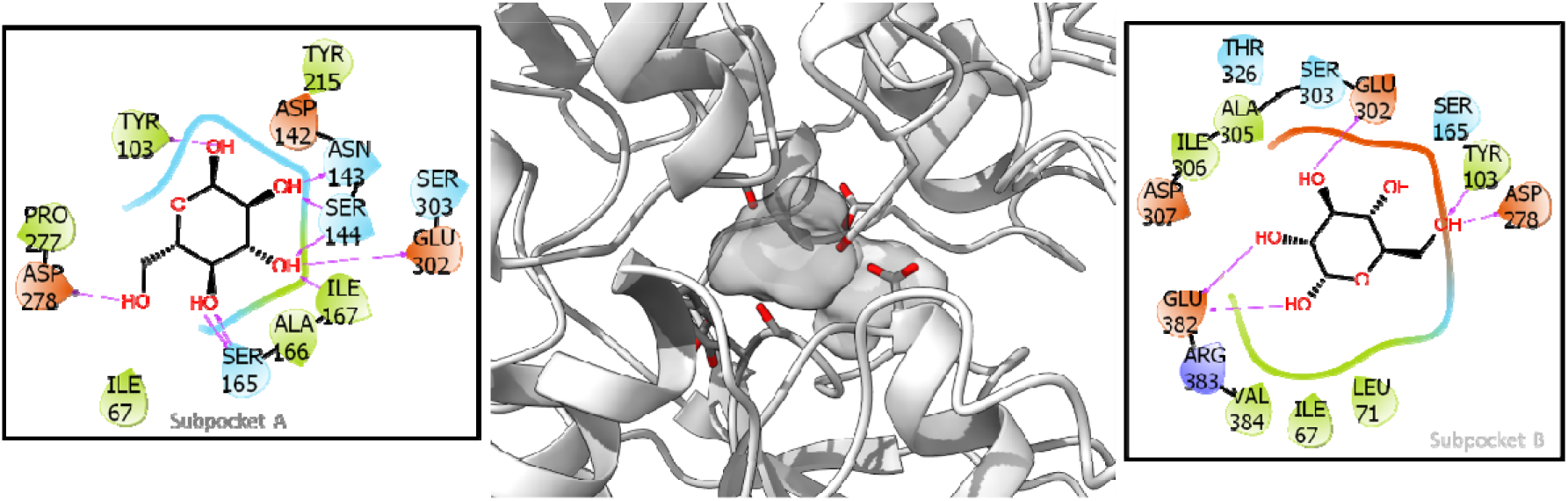
Subpocket A is shown in dark gray and also in 2D in the left panel (with docked l-glucose). Subpocket B is shown in light gray and also in 2D in the right panel (with docked d-glucose).

Other glucose-binding proteins are known to be stereospecific. For example, the enzyme hexokinase acts on d-glucose and not on l-glucose (Aleshin et al., 1998). We docked l-glucose into the active site instead of the co-complexed d-glucose found in the crystal structure (PDB ID: 1HKB). Unlike in Tas1R2, docking of l-glucose in the d-glucose-binding site in hexokinase is not possible without its rearrangement (Figure S3). Glucose transporter 1 (GluT1) is another example of stereospecificity toward glucose. Previous work (Cunningham, Afzal-Ahmed, & Naftalin, 2006) found two stereospecific binding sites that could bind d-glucose but not l-glucose. This was attributed to the 3-amino acid motif QLS (not found in Tas1R2).

## Discussion

In this study, we assessed whether l-glucose elicits sweet taste via taste receptor Tas1R2/Tas1R3. We confirmed that l-glucose is perceived as sweet, showing that sweetness ranking of d- and l-glucose by individuals varies only slightly and is similar on average.

We next showed that sweetness of l-glucose is mediated by Tas1R2/Tas1R3 sweet taste receptor using transiently transfected HEK293T cells. In cells, stronger activation of d-glucose compared to l-glucose was observed. Differences between in-vivo taste perception and in-vitro taste receptor activation are common, and have been reported in other studies (Cheled-Shoval et al., 2017). The difference can stem from several reasons, including variation in exposure time to the stimulus, unequal expression levels in the human palate compared to transfected cells, and sensory properties and interactions of tastants with saliva or volatile flavor elements (in in-vivo systems) compared to pure substances (in in-vitro assays).

Computational analysis, including d- and l-glucose docking to the homology model of the Tas1R2 VFT domain, identified two glucose-binding sites. One is close to the hinge between the two lobes, its binding residues described for aspartame and neotame (Liu et al., 2011; Maillet et al., 2015). Subpocket B, adjacent to subpocket A, overlapped with that which has been described for sweetness enhancers (F. Zhang et al., 2010). The residues defining each of the subpockets are conserved among primates and rodents, as expected from their ability to sense natural sugars. The exceptions are I67 and D278, which are replaced by residues with similar properties. Indeed, similar sweetness of l-glucose compared to d-glucose for mice was demonstrated by the same licking rates of d- and l-glucose solutions. This indicated similar sweetness of the enantiomers for mice (Tellez et al., 2016) in accordance with the in-vivo results in humans.

The two subpockets of the d- and l-glucose enantiomers may potentially be used by combinations of sweet compounds. Such combinations were previously shown to have enhancing effects on sweetness perception (S. S. Schiffman, Sattely-Miller, Graham, Booth, & Gibes, 2000),(Keast & Breslin, 2003). Specifically, aspratame and d-glucose mixture is significantly sweeter than the average of two pure solutions (Gliemmo, Calviño, Tamasi, Gerschenson, & Campos, 2008; S. S. Schiffman et al., 1995). Our findings suggest a combined structural and functional assay-based framework for future rational design and optimization of sweeteners combinations.

The comparison to other stereo-specific glucose binding proteins showed that hexokinase active site could not contain l-glucose without a steric clash. Additionally, previous work that discussed binding of d- and l-glucose to GluT1 (Cunningham, Afzal-Ahmed, & Naftalin, 2006) found two stereo-specific binding sites that could bind d-glucose but not l-glucose. This was attributed to the 3-amino acid motif QLS. The QLS motif was found to be unique to GluT1 but not to Tas1R2. Thus, glucose has different types of pockets in its different targets, some stereospecific and others not.

Our integrated in-vivo, in-vitro and in-silico study provides a suggestion of glucose enantiomers binding modes in sweet taste receptors, and may offer future directions for sweeteners combinations, targeting the two subpockets.

## Materials and Methods

### In-vivo (sensory) experiments

The 2AFC experiments aimed to evaluate the sweetness of sucrose vs. d-glucose and of d-glucose vs. l-glucose: 27 participants were asked to choose the sweeter sample out of two alternating 0.48 M sugar solutions (volume of 0.3 mL).

Dose-response to d-glucose and l-glucose sweetness: 11 healthy, 21- to 40-year-old panelists were trained with five concentrations of sucrose solution (0.15, 0.24, 0.48, 0.96 and 1.5 M) and three concentrations of d-glucose solution (0.24, 0.48 and 0.96 M). Volunteers who were smokers, pregnant, or had specific chronic diseases or medical conditions were excluded from participation in the experiments. The results were analyzed and the most consistent trainees (n = 6) participated in the subsequent experiments, following (Barry G. Green, Shaffer, & Gilmore, 1993): these six participants used the ‘sip & spit’ method and the gLMS to evaluate the sweetness of d- and l-glucose (B. G. Green et al., 1996). The taste experiments were prepared by Compusense cloud (www.compusense.com, Rev-11/05/2017). The gLMS is a semantic scale of perceptual intensity characterized by quasi-logarithmic spacing of its verbal labels. It encompasses a wide numerical range between its lowest (‘barely detectable’) and highest (‘strongest imaginable’) verbal descriptors. Statistical analyses were carried out in the JMP Pro statistical software package. Data were analyzed by paired *t*-test, with an alpha value of 0.05 (version 13.0.0; SAS Institute Inc.).

The sensory tests took place in a designated room with minimal distraction for the participants. An institutional review board ethics committee approved the experiments; each participant signed a consent form and was paid for their participation in these experiments. The experiments were blinded—the participants were not aware of the tested compounds’ order or content, each sample being tagged with a 3-digit code. Each experiment had a specific total glucose concentration, and varying percentage substitution with l-glucose (from zero up). The participants’ ages varied from 21 to 40 years. All of the participants were instructed to arrive on the same day, and to refrain from coffee or a heavy meal for 1 h prior to the experiment. Upon arrival, each participant signed the consent form and was handed an instructional page on how to use the gLMS.

Ratings of intensity on the gLMS are relative to the ‘strongest imaginable’ oral sensation of any kind. In addition, each participant was briefed by one of the experimenters to clarify the use of the gLMS and to answer any questions. The experiment was divided into sessions with different concentrations (in increasing order), with a 5-min break between sessions. In each session, the recipients were given aliquots of samples with substituted glucose by percentage. The volumes were 0.3 mL (dripped with a sterile syringe on the center of the tongue by the experimenters) or 3 mL (sip & spit). The participants were asked to rate the sweetness intensity on the gLMS presented on a computer screen. After each sample, the participants rinsed their mouth with water and spit it out, and were instructed to wait for at least 30 s before advancing to the next sample.

### Computational analysis

#### Modeling

The sweet taste receptor sequence was obtained from the Uniprot database (hT1R2, ID: Q8TE23). The I-Tasser server was used to create models of the hT1R2 VFT using the specified template PDB 5X2M chain B. The model was minimized and prepared for docking with the Protein Preparation Wizard tool in Maestro and Glide Grid Generation (Schrodinger tools 2019-1).

#### Ligand preparation

The ligands were prepared for docking using LigPrep. Conformers, tautomers and protomers (different protonation states of the ligands) were enumerated at pH 7.0 ± 1.0, retaining the specified chiral centers (Schrodinger tools 2019-1).

#### Docking protocol

The binding site was defined as a 12-Å grid around the l-glutamate-binding site in the class C GPCR mGluR1 (PDB ID: 1EWK). Overlap of the models with the crystal structure was used to define the binding grid in the model. The docking protocol included Maestro Schrodinger 2019-1, Glide Extra Precision mode (XP), flexible ligand sampling, and Glide XP docking scores.

#### IFD

We performed IFD for each ligand after docking with the initial XP docking protocol. After docking of α or β, d- or l-glucose (Figure 9), the same compound was docked over the initial docking of the initial ligand. That is, α-d-glucose induced-fit complexes were predicted from a starting complex of a docked α-d-glucose, ending with 11 different complexes. The same was applied for β-d-glucose (18 IFD complexes), α-l-glucose (6 IFD complexes) and β-l-glucose (15 IFD complexes). Each molecule was docked via IFD based on a starting pose from each relevant monosaccharide, as performed in a previously published protocol used with the same model (Ben Shoshan-Galeczki & Niv, 2020).

**Figure 9.**
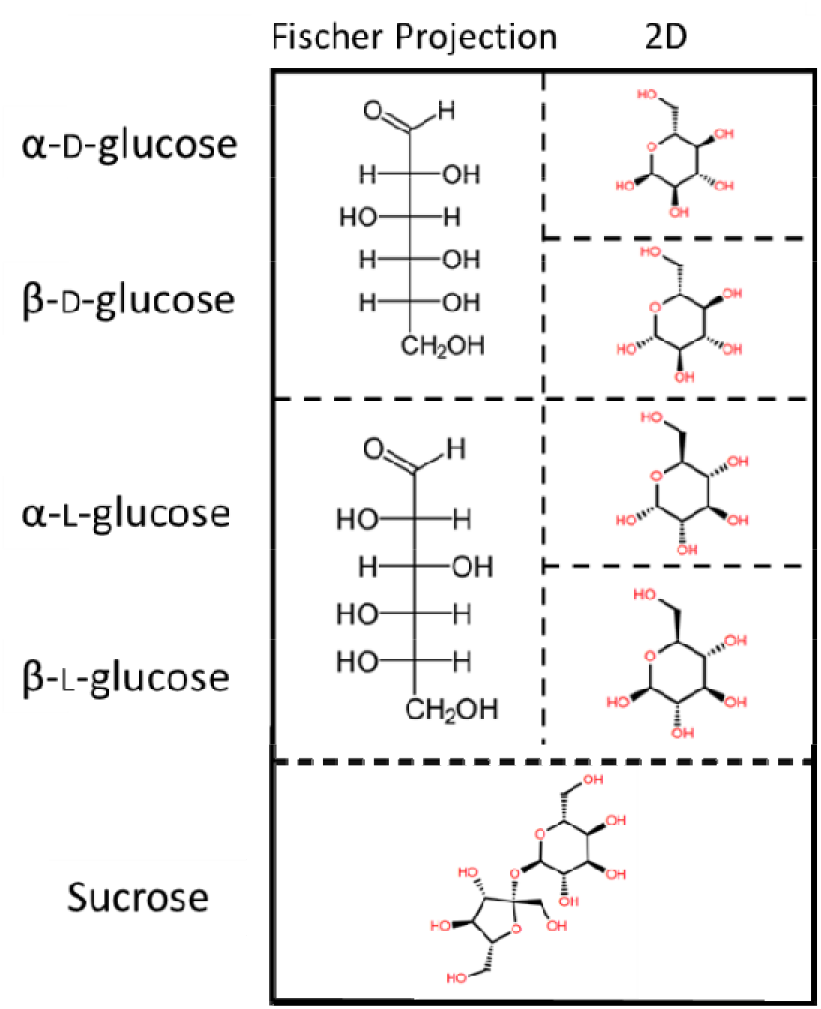
The most common isomers of d- and l-glucose in Fischer projection and 2D representation. Sucrose is shown in 2D representation.

### Cell-based system

#### Materials

d-glucose, l-glucose, sucrose, sucralose, acesulfame-K, Dulbecco’s Modified Eagle’s medium (DMEM) and lithium chloride (LiCl) were purchased from Sigma. Lipofectamine™ 2000 was purchased from Invitrogen.

#### Cell culture

HEK293T cells, obtained from the American Type Culture Collection (ATCC), were maintained in 10% (v/v) DMEM supplemented with 10% (v/v) fetal bovine serum (FBS), 1% (v/v) l-glutamine amino acid, and 0.2% (v/v) penicillin–streptomycin. Cells were kept in the incubator at 37 °C, in a humidified atmosphere containing 5% CO_2_.

#### IP1 assay

Transfection was performed on a 10-cm plate. The desired plasmids (TAS1R2, TAS1R3, Gα16gus44) were transiently transfected into HEK293T cells using Lipofectamine™ 2000. One day after transfection, transfected cells were suspended in 10% DMEM, and seeded onto a 24-well culture plate coated with poly-d-lysine. The plate was then kept in the incubator for 8 h to obtain cell adherence. Cells were then “starved” by changing the medium to 0.1% DMEM (containing 0.1% FBS) to reduce their basal activity. After 16–18 h of starvation, cells were exposed to the ligands dissolved in 0.05 M LiCl, because LiCl is crucial for IP1 accumulation. Solution pH was 7.4 (+0.5). Cells were exposed to the tastants by adding the tastant solution directly into the wells, for 5 min. At the end of the exposure time, the well content was replaced with fresh 0.1% DMEM containing LiCl, for another 55-min incubation. At the end of the incubation, the wells were washed with phosphate buffered saline (PBS) + Triton X-100 1% (v/v), and kept at −80 ⍰C to disrupt the cell membrane. The cell content was collected and used for IP1 assay (the homogeneous time-resolved fluorescence IP1 assay was used to measure intracellular IP1 accumulation according to the manufacturer’s directions (Cisbio Bioassays, Perkin Elmer). In brief, 15μl cell lysate was added to each well of a 384-well plate, with the addition of IP1-d2 conjugate and IP1 monoclonal antibody (dissolved in lysis buffer). After 1-h incubation at room temperature, the plate was read in a microplate reader (CLARIOstar, BMG Labtech). The resulting signal was inversely proportional to the concentration of IP1 in the sample. Dose-response curves were fitted by Prism 8 (Graphpad).

## Acknowledgements

We thank Dr. Tzafi Danieli for help and advice with the cell-culture and transfection. We thank Dr. R. F. Margolskee for the pcDNA3 of chimeric Gα16gust44 and Dr. Maik Behrens for the pcDNA3 of T1R3, and for the pcDNA5FRT PM of T1R2. Nitzan Dubovski is a recipient of the Robert H. Smith Fellowship for Excellence in Agriculture, Food and Environment. The study is supported by ISF grant #1129/19 and UHJ-France and the Foundation Scopus.

