## SupplementarySweetChirality for "Sweet chirality: the taste of l- and d-glucose stereoisomers"

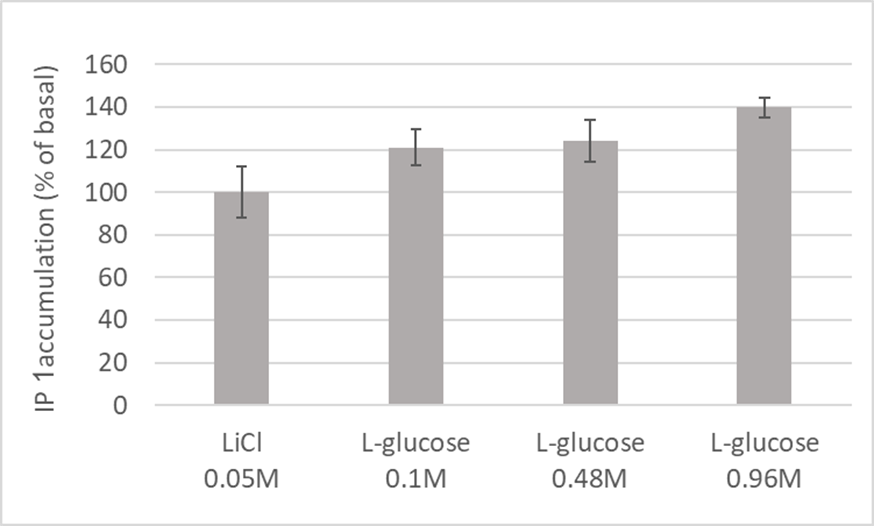

**Figure S1:** Exposure of non-transfected HEK293T cells to L-glucose concentration has evoked no significant response, compared to basal level (0.05M LiCl).

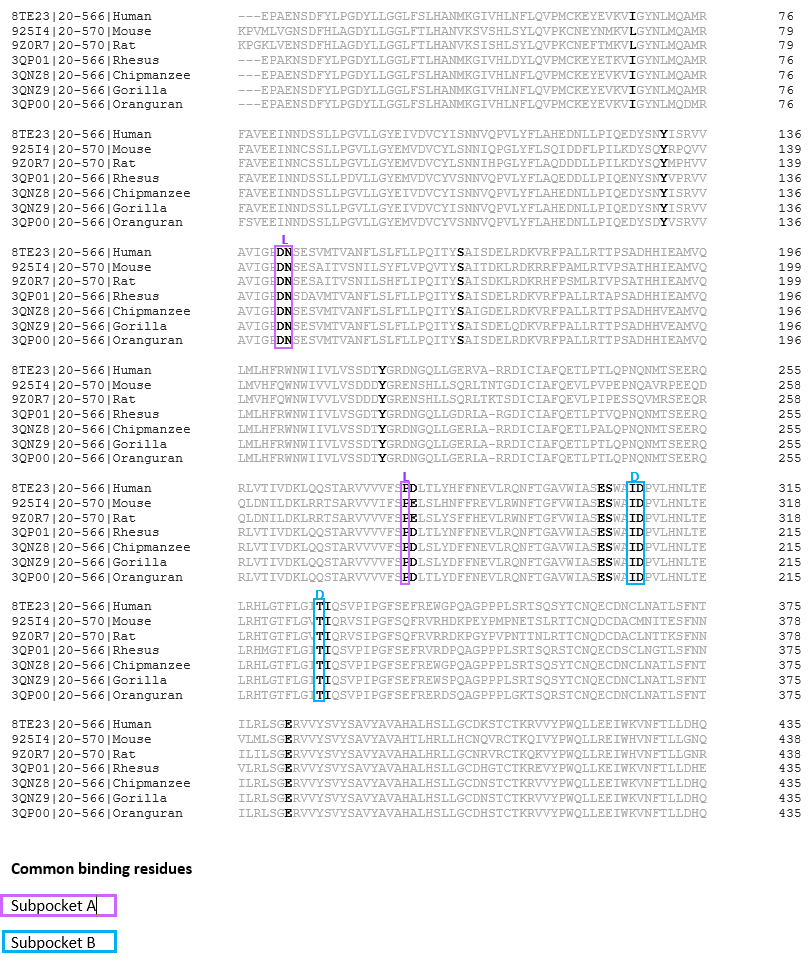

**Figure S2:** Multiple sequence alignment of T1R2 VFTM between different mammals. The Uniprot accession code is annotated in the beginning of each sequence. Residues that were involved in binding of D- or L-glucose are colored in black. Residues that were involved in binding only in the first subpocket are highlighted in purple and residues that were involved in binding in the second subpocket are highlighted in cyan. Residues that are highlighted in black were involved in both subpockets.

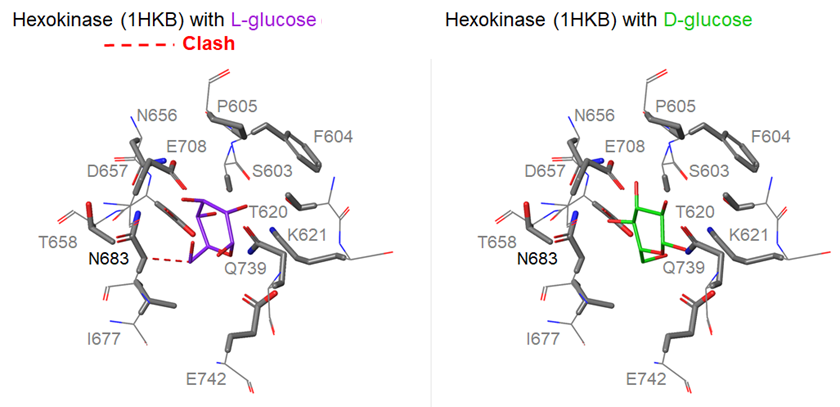

**Figure S3:** Hexokinase binding site after minimization for D- and L-glucose. The figure is based on the 1HKB (Aleshin et al., 1998) crystal structure binding site.
